# Pain and breathlessness: Salient, somatosensory and similar, but not the same

**DOI:** 10.1101/2020.05.04.076174

**Authors:** Olivia K. Harrison, Anja Hayen, Tor D. Wager, Kyle T. S. Pattinson

## Abstract

Quantifying pain currently relies upon subjective self-report. Alongside the inherent variability embedded within these metrics, added complications include the influence of ambiguous or prolonged noxious inputs, or in situations when communication may be compromised. As such, there is continued interest in the development of brain biomarkers of pain, such as in the form of neural ‘signatures’ of brain activity. However, issues pertaining to pain-related specificity remain, and by understanding the current limits of these signatures we can both progress their development and investigate the potentially generalizable properties of pain to other salient and/or somatomotor tasks. Here, we utilized two independent datasets to test one of the established Neural Pain Signatures (the NPS (Wager et al. 2013)). In Study 1, brain activity was measured using functional magnetic resonance imaging (fMRI) in 40 healthy subjects during experimentally induced breathlessness, conditioned anticipation of breathlessness and a simple finger opposition task. In Study 2, brain activity was again measured during anticipation and breathlessness in 19 healthy subjects, as well as a modulation with the opioid remifentanil. We were able to identify significant NPS-related brain activity during anticipation and perception of breathlessness, as well as during finger opposition using the global NPS. Furthermore, localised NPS responses were found in early somatomotor regions, bilateral insula and dorsal anterior cingulate for breathlessness and finger opposition. In contrast, no conditions were able to activate the local signature in the dorsal posterior insula - thought to be critical for pain perception. These results provide properties of the present boundaries of the NPS, and offer insight into the overlap between breathlessness and somatomotor conditions with pain.

## Introduction

Whilst perceptions of pain are often identified and assessed through subjective self-report, these experiences are influenced by higher cognitive functions such as attention (Wiech et al. 2008) and expectation (Atlas & Wager 2012). Furthermore, pain perception can be altered with prolonged noxious inputs, and are potentially difficult to quantify in infants, in those who have cognitive impairment, or those who are minimally conscious (Wager et al. 2013). Therefore, the quest has begun for biological ‘readouts’ related to pain in the brain, with the hope of allowing us to assess pain within an individual using non-invasive neuroimaging measures (Wager et al. 2013; Woo, Schmidt, et al. 2017). These tools are designed to identify pain across experiments and laboratories, and eventually lead to use in those who cannot accurately express pain for themselves.

Here we focus on the Neurologic Pain Signature (NPS), an established pain-related brain measure. An advantage of this measure is that it has been widely tested—on over 40 unique participant cohorts to date—for sensitivity and specificity to pain, generalizability across populations and evoked pain types, and other properties (for reviews, see (Woo, Chang, et al. 2017; Kragel et al. 2018)). The NPS is a distributed pattern of activity across brain regions, including the major targets of ascending nociceptive pathways (dorsal posterior insula, ventrolateral and medial thalamus, mid- and anterior insula, anterior midcingulate, amgydala, periaqueductal gray, hypothalamus). It can be applied to individual-person level data across studies (Wager et al. 2013), yielding an objective brain measure (Woo & Wager 2016)1. Applying the NPS entails calculating a weighted average across voxels for a test functional brain image (i.e., the dot product) or another pattern similarity metric. Pattern weights limited to individual regions can also be used to obtain local pattern responses (Woo et al. 2014).

This approach is part of a major trend in neuroimaging research using pattern information to assess pain (Rosa & Seymour 2014; Mano et al. 2018; van der Miesen et al. 2019; Ung et al. 2012; Marquand et al. 2010) and other cognitive and affective processes. Multivariate brain models integrate brain information into a single optimized prediction, and test predictions on new, independent individuals, providing unbiased estimates of effect size (Reddan et al. 2017) and capturing information across multiple spatial scales (Miyawaki et al. 2008; Hackmack et al. 2012; Haynes 2015; Lindquist et al. 2017). NPS responses have also been found to correlate with the intensity of variations in evoked experimental pain in individuals across multiple studies (Lindquist et al. 2017; Woo, Schmidt, et al. 2017). In one analysis across 6 studies (N = 180), NPS responses were positively correlated with trial-by-trial pain reports in 93% of individual participants (Lindquist et al. 2017). The NPS has also been shown to demonstrate some specificity towards somatic pain: It does not respond to non-noxious warm stimuli (Wager et al. 2013), threat cues (Wager et al. 2013; Krishnan et al. 2016; Ma et al. 2016), social rejection-related stimuli (Wager et al. 2013), observed pain (Krishnan et al. 2016), or aversive images (Chang et al. 2015), although many of these conditions are affective, salient, and activate many of the same gross anatomical regions as somatic pain. Therefore, the NPS is not a complete model for all types of and influences on pain (Woo, Schmidt, et al. 2017), but rather appears to track pain of nociceptive origin (including thermal, mechanical, laser, visceral, and electrical; (Krishnan et al. 2016; Woo & Wager 2016; López-Solà et al. 2017; Zunhammer et al. 2018)) in a fashion that is relatively insensitive to cognitive input. It does not respond to social ‘pain’ (Woo et al. 2014; Krishnan et al. 2016), and it is not strongly influenced by placebo treatment (Zunhammer et al. 2018), cognitive regulation (Woo et al. 2015), reward (Becker et al. 2017), knowledge about drug-delivery context (Wager et al. 2013; Zunhammer et al. 2018), or perceived control (Bräscher et al. 2016). On the other hand, the NPS does show significant responses to remifentanil, citalopram, spinal manipulation (in chronic neck pain suffers), and some types of psychosocial/behavioral manipulations, showing promise as a pharmacodynamic biomarker. These findings underscore the idea that the NPS and other brain measures do not “measure pain” (a subjective experience), but rather measure specific neurophysiological processes linked to pain construction.

However, whilst early results have proven promising when delineating pain from other emotion-based stimuli, these measures have not typically been tested against predominantly somatosensory aversive stimuli. One ideal test case might be the frightening perception of breathlessness; a multi-dimensional symptom that causes major suffering across a broad range of individuals (Marlow et al. 2019; Hayen et al. 2013; Herigstad et al. 2011). In fact, the definition of breathlessness (or ‘dyspnea’) from the American Thoracic Society draws many comparisons that closely parallel perceptions of pain (Parshall et al. 2012), and previous work has noted many similarities between brain networks associated with both breathlessness and pain (Leupoldt et al. 2009). However, whether this broad correspondence is represented within more highly localized pain signatures, and what this means for our understanding of these vastly different perceptions, is not yet known. Furthermore, isolated somatomotor activity has also yet to be exclusively tested against these pain signatures, many of which load heavily on somatomotor networks within the brain (Cauda et al. 2012).

Here, we aimed to test the specificity of the Neural Pain Signature (NPS, (Wager et al. 2013)) using salient and somatomotor tasks. We employed two datasets that induced both the anticipation and perception of breathlessness (Study 1 – collected at 7 Tesla in 40 healthy subjects (Faull & Pattinson 2017); and Study 2 – collected at 3 Tesla in 19 healthy subjects (Hayen et al. 2017)), and a simple somatomotor task of finger opposition (Study 1). We then investigated local patterns of pain-related activity from the regional NPS responses, allowing us to disentangle where the major similarities or differences may exist between these conditions. Additionally, we explored the effect of opioid administration (Study 2) to test the potential modulation of the global and regional NPS responses to both the anticipation and perception of breathlessness. We aimed to find the boundary conditions for the NPS to both understand existing limitations and generalizable properties as a biomarker for pain, support its refinement towards greater pain specificity, and investigate the potential neural similarities and differences between pain and breathlessness.

## Methods

### Testing data sets

To test the current limitations of the NPS, data from previously published work was utilized in these analyses (Faull & Pattinson 2017; Hayen et al. 2017) (please see previous publications for a full description of the study methods, scanning protocols and univariate analyses). Briefly, the first dataset was acquired at 7 Tesla (Faull & Pattinson 2017), and employed one level of breathlessness (induced by inspiratory resistive loading) during fMRI, with preceding anticipation periods cued by conditioned shapes presented on the screen. Control tasks of no anticipation or breathlessness (cued via the presentation of a conditioned shape that was never paired with breathlessness) and finger opposition (cued by the word ‘tap’ presented on the screen) were also collected. Each condition was presented 14 times in a pseudo-randomised order. The contrasts of interest that were analysed against the NPS for this study were anticipation > no breathlessness cue (‘Anticipation’ contrast), breathlessness > no breathlessness (‘Breathlessness’ contrast), and finger opposition > baseline (‘Finger opposition’ contrast).

The second dataset was acquired at 3 Tesla (Hayen et al. 2017), and employed two levels of breathlessness (mild and strong, also induced with inspiratory resistive loading) with conditioned anticipation periods, and a cued control condition of no anticipation or breathlessness (as above). Four repeats of each of the paired anticipation and breathlessness cues were presented, and eight repeats of the unloaded condition were performed (pseudo-randomised order). This study involved two scans, with either a controlled infusion of the opioid remifentanil (0.7 ng/ml target) or saline placebo (single-blind, counterbalanced order). For this analysis, we have not considered the anticipation and perception of mild breathlessness, to remain consistent and attempt to replicate any results found in Study 1. The contrasts of interest that were analysed against the NPS here were anticipation of strong breathlessness > no breathlessness cue in the saline condition (‘Anticipation’ contrast), strong breathlessness > no breathlessness in the saline condition (‘Breathlessness’ contrast), anticipation of strong breathlessness > no breathlessness cue in the remifentanil condition (‘Remi Anticipation’ contrast), and strong breathlessness > no breathlessness in the remifentanil condition (‘Remi Breathlessness’ contrast). The difference between saline and remifentanil conditions were also compared for both anticipation and breathlessness contrasts.

### NPS analyses

For each contrast in each study, we calculated the overall NPS response as specified by Wager and colleagues (Wager et al. 2013). This entailed taking the dot product of the NPS weight map and each test contrast image from each individual participant, calculating a weighted average over each test image, where the NPS map specifies the weights. It reduces each contrast image to a single number, the ‘NPS response’, which is the predicted pain intensity based on the model. We tested whether the NPS responses were significantly different from zero using standard t-tests. This is mathematically equivalent to conducting paired t-tests on within-person contrasts, treating participant as a random effect. We also applied the local NPS patterns from nociceptive target regions with predominantly positive weights (‘NPS Positive’ subregions) and regions with negative weights (‘NPS Negative’ subregions), as defined in (López-Solà et al. 2017) and (Krishnan et al. 2016). We use a standard threshold of p < 0.05 for statistical significance in these a priori tests (one star in figures), and also note tests that are significant at p < 0.01 (two stars) and q < 0.05 False Discovery Rate corrected (three stars). Finally, we tested whether NPS responses were related to sedation levels and the order in which conditions were administered.

## Results

Anticipation of breathlessness, breathlessness perception and finger opposition all significantly activated the overall NPS (Table 1 and Figure 1), and the findings for anticipation and breathlessness were able to be replicated in two independent datasets. The administration of remifentanil in Study 2 did not alter the NPS response to anticipation of breathlessness, and while it appeared to reduce the response to breathlessness itself, this did not reach statistical significance (Table 1).

**Table 1.**
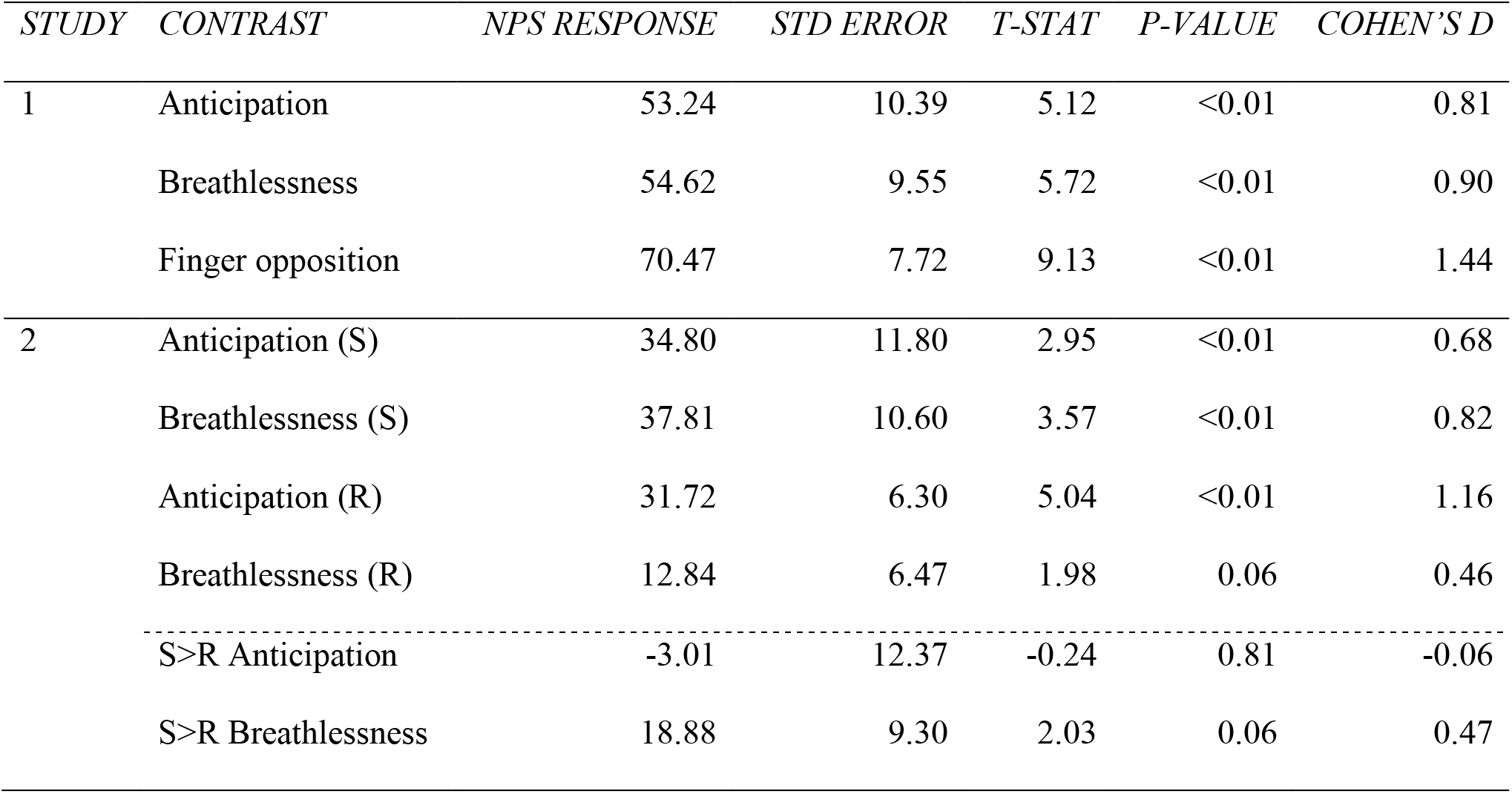
NPS responses and statistics for the contrasts of interest in each study. Study 1 was conducted at 7 Tesla with 40 participants and 14 stimulus repeats, while Study 2 was collected at 3 Tesla with 19 participants and 4 stimulus repeats.

**Figure 1.**
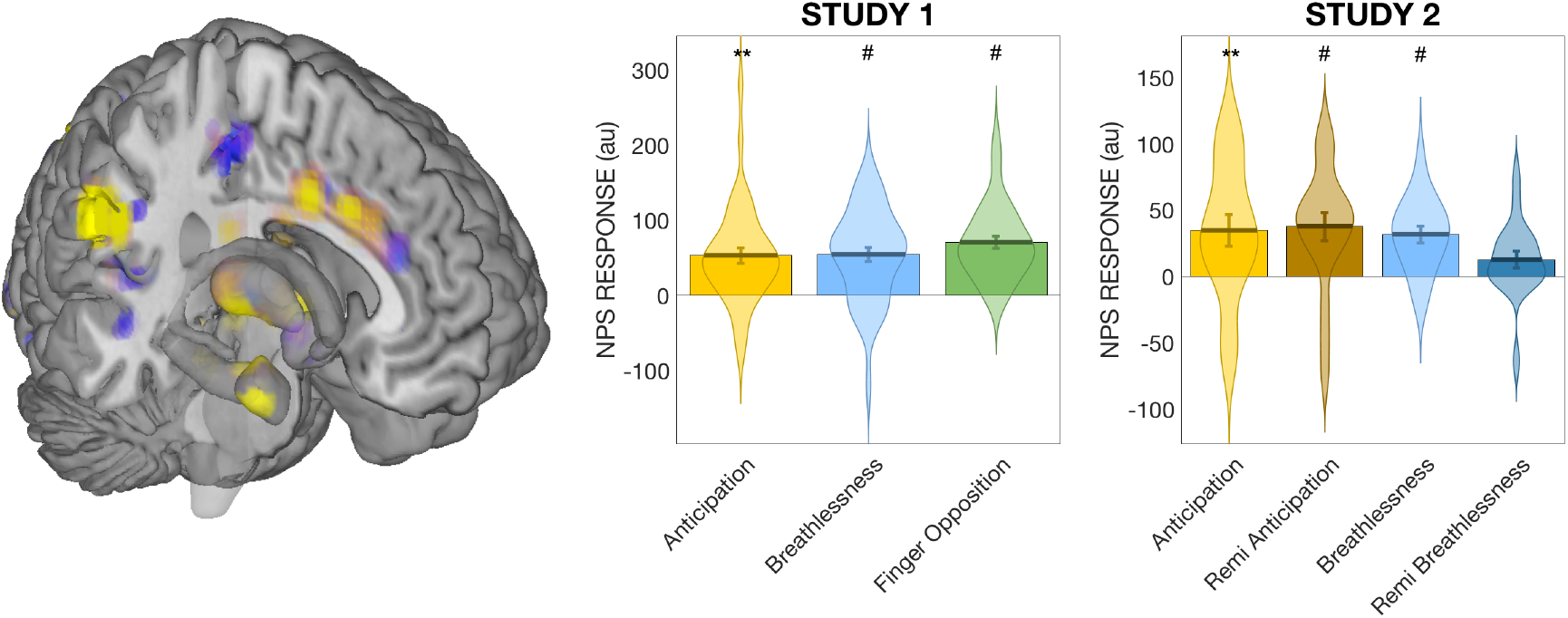
Overall NPS activity in the contrasts of interest for the two datasets. Left: Three-dimensional representation of some of the core regions of the NPS. ** Significantly different from zero at p < 0.01 (most satisfy p < 0.001); # Significantly different from zero at q < 0.05 (FDR corrected).

### Study 1 regional NPS results

Within the NPS subregions, the anticipation contrast produced significant responses in the positive NPS regions of the bilateral insula, and significant responses in the negative NPS regions of the bilateral lateral occipital cortex and right inferior parietal lobule (Figures 2 and 3; Supplementary Table 1). During breathlessness, significant responses were observed in the positive NPS regions of the bilateral insula, right thalamus, right secondary sensory cortex, dorsal anterior cingulate cortex and vermis, and significant responses in the negative NPS region of the right inferior parietal lobule (Figures 2 and 3; Supplementary Table 1). Consistent with the breathlessness contrast, finger opposition also produced significant responses in the positive NPS regions of the bilateral insula, right thalamus, right secondary sensory cortex, dorsal anterior cingulate cortex and vermis, plus additional activity in the right primary visual cortex. In the negative NPS regions, finger opposition activated the lateral occipital cortex and right posterior lateral occipital cortex (Supplementary Table 1). No contrasts produced significant activity in the right dorsal posterior insula subregion of the NPS (Figure 2). Full statistical reports and visualisations of the raw condition-related activity are provided in the supplementary material.

**Figure 2.**
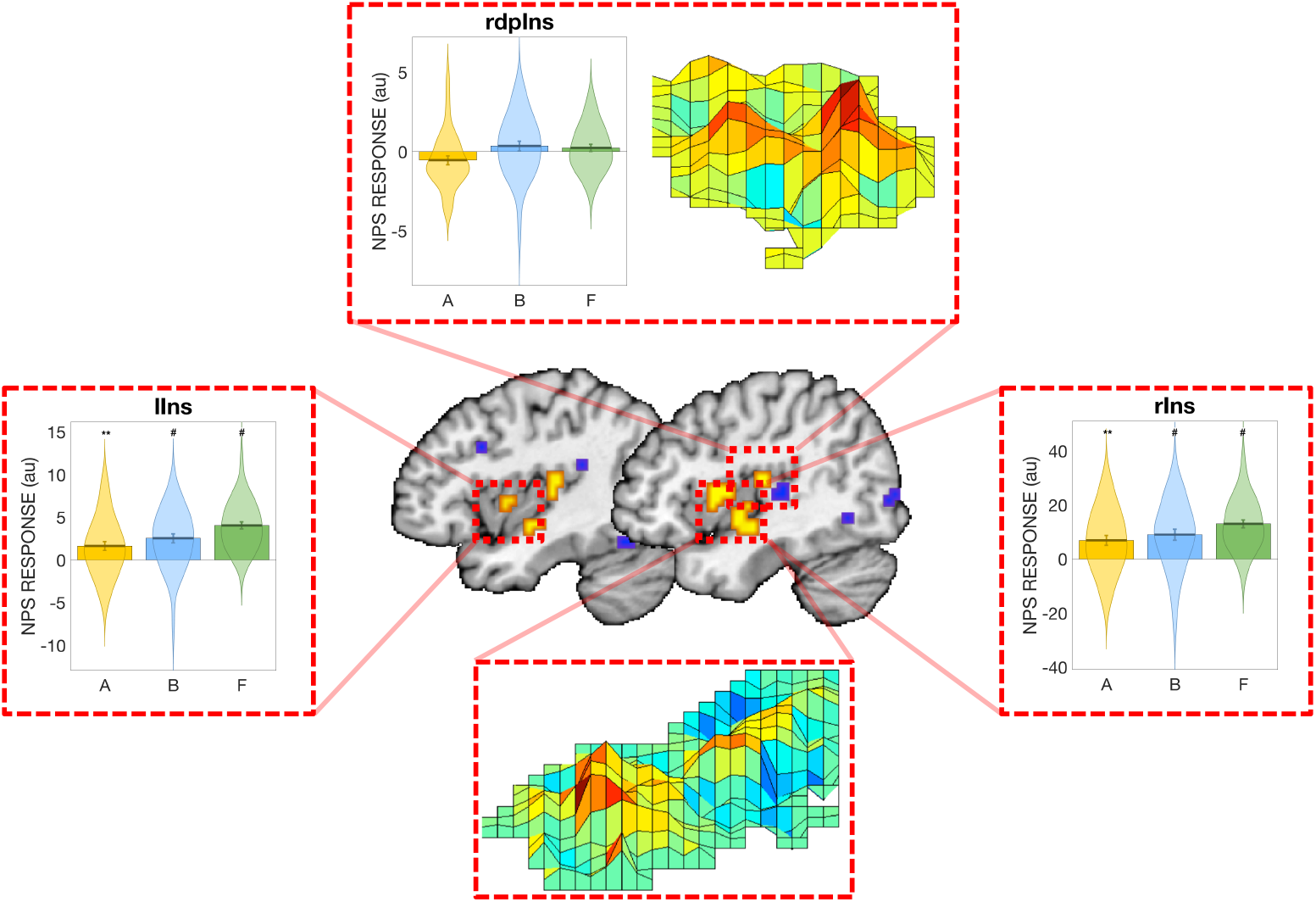
Regional NPS activity in the insula for the anticipation, breathlessness and finger opposition contrasts from Study 1. Robust statistical activity is observed in the bilateral insula (labelled lIns and rIns) for all three conditions, while no significant activity is observed in the right dorsal posterior insula (rdpIns). Abbreviations: A, Anticipation contrast; B, Breathlessness contrast; F, Finger opposition contrast. ** Significantly different from zero at p < 0.01; # Significantly different from zero at q < 0.05 (FDR corrected).

**Figure 3.**
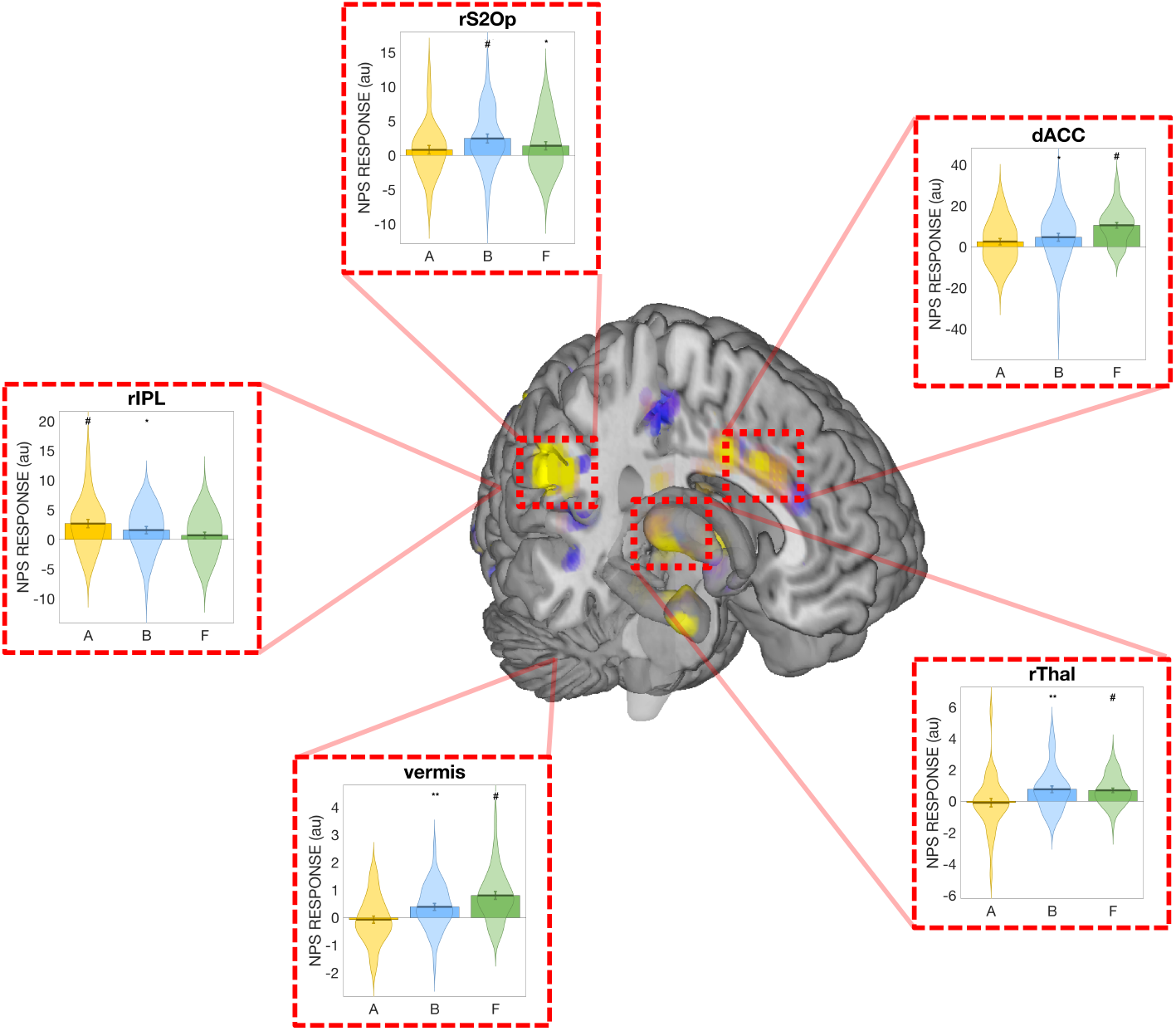
Regional NPS activity subregions of the NPS for the anticipation, breathlessness and finger opposition contrasts from Study 1. Significant NPS activation is observed in the dorsal anterior cingulate cortex (dACC), right thalamus (rThal), right secondary somatosensory cortex / operculum (rS2Op) and vermis for both breathlessness and finger opposition, and in the right inferior parietal lobule (rIPL) for both anticipation and breathlessness. For a full list of regions please see Supplementary Table 1. Abbreviations: A, Anticipation contrast; B, Breathlessness contrast; F, Finger opposition contrast. * Significantly different from zero at p < 0.05; ** Significantly different from zero at p < 0.01; # Significantly different from zero at q < 0.05 (FDR corrected).

### Study 2 regional NPS results

Within the positive NPS subregions in Study 2, the anticipation (saline) contrast produced a significant response in the right primary visual cortex, with a negative response in the right dorsal posterior insula (Figures 4 and 5; Supplementary Table 2). No significant responses were found in the negative NPS subregions. The administration of remifentanil did not significantly modulate any of the NPS-related subregion activity during anticipation, although the right insula (positive region) and right posterior lateral occipital cortex and left superior temporal sulcus (negative regions) all additionally produced significant results (Figures 4 and 5; Supplementary Table 2).

**Figure 4.**
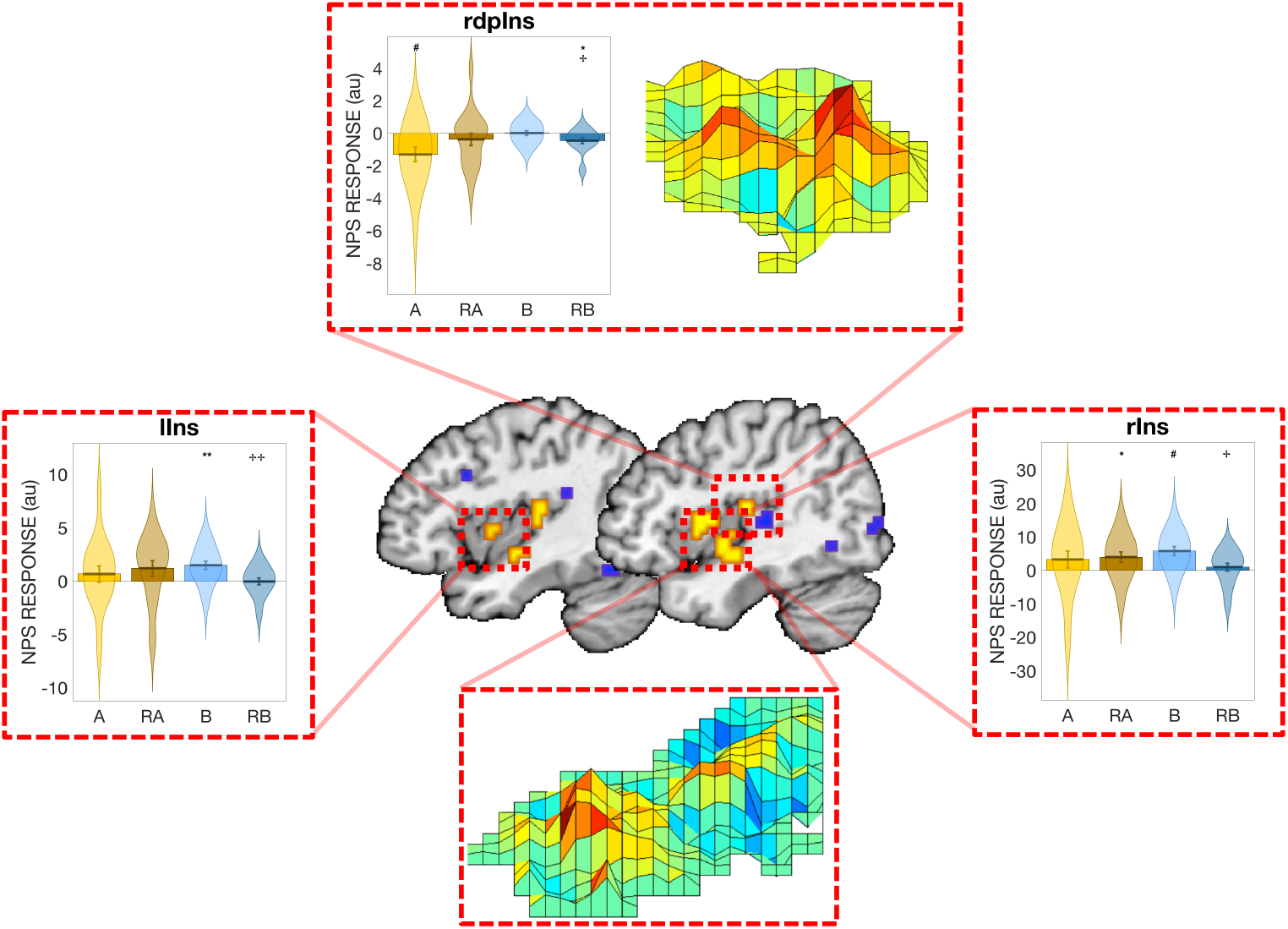
Regional NPS activity in the insula for the anticipation and breathlessness contrasts during both saline and remifentanil administration from Study 2. Robust, positive statistically significant NPS-related activity is only observed in the bilateral insula (labelled lIns and rIns) for the breathlessness condition, which is significantly modulated by the administration of the opioid remifentanil. NPS-related activity in the right dorsal posterior insula (rdpIns) is significantly decreased during saline anticipation. Abbreviations: A, Anticipation contrast (saline); RA, Remifentanil anticipation contrast; B, Breathlessness contrast (saline); RB, Remifentanil breathlessness contrast. ** Significantly different from zero at p < 0.01; # Significantly different from zero at q < 0.05 (FDR corrected); ^✢✢^ Significantly modulated by remifentanil at p < 0.05.

**Figure 5.**
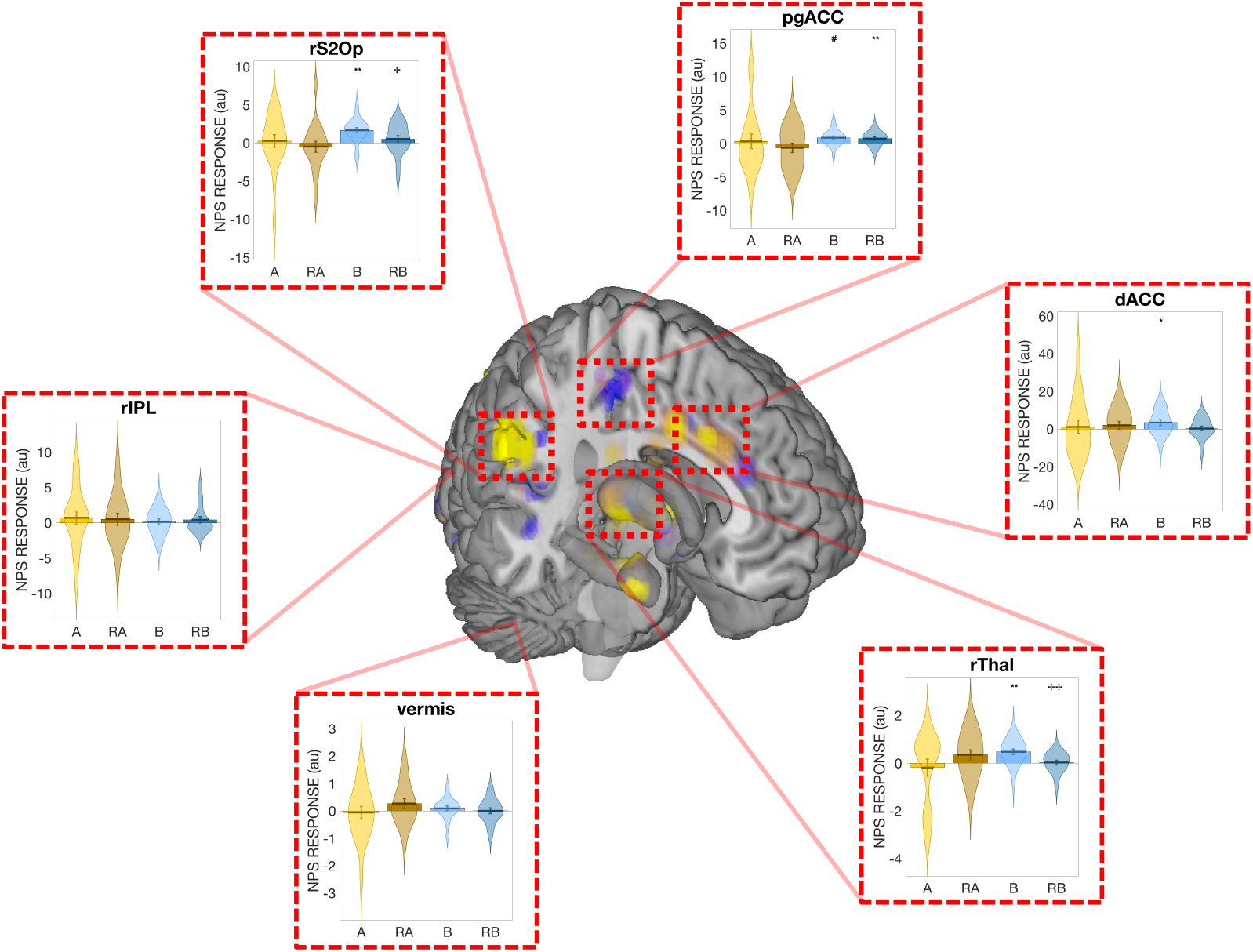
Regional NPS activity subregions of the NPS for the anticipation and breathlessness contrasts during both saline and remifentanil administration from Study 2. Significant NPS activation is observed in the dorsal and pregenual anterior cingulate cortex (dACC and pgACC), right thalamus (rThal) and right secondary somatosensory cortex / operculum (rS2Op) for breathlessness, with the NPS-related activity in the right thalamus and rS2Op significantly modulated by the administration of the opioid remifentanil. For a full list of regions please see Supplementary Table 2. Abbreviations: A, Anticipation contrast (saline); RA, Remifentanil anticipation contrast; B, Breathlessness contrast (saline); RB, Remifentanil breathlessness contrast. * Significantly different from zero at p < 0.05; ** Significantly different from zero at p < 0.01; # Significantly different from zero at q < 0.05 (FDR corrected); ^✢^ Significantly modulated by remifentanil with p < 0.05; ^✢✢^ Significantly modulated by remifentanil at p < 0.01.

During breathlessness, the positive NPS regions of bilateral insula, right thalamus, right secondary sensory cortex and dorsal anterior cingulate cortex produced significant NPS-related activity, while the negative NPS subregion of the pregenual anterior cingulate cortex was also significant (Figures 4 and 5; Supplementary Table 2). The administration of remifentanil significantly decreased the NPS-related activity in all saline significant regions except the pregenual anterior cingulate cortex, and additionally produced a significant decrease in the right dorsal posterior insula (Figures 4 and 5; Supplementary Table 2).

## Discussion

### Main findings

Utilising two independent datasets, we have demonstrated that both the anticipation and perception of breathlessness robustly evoked significant activity in an established pain signature (NPS (Wager et al. 2013)), and this NPS-related activity during breathlessness was able to be modulated by the infusion of the short-acting opioid remifentanil (Study 2). Furthermore, a somatomotor finger opposition task was also able to evoke significant activity within the NPS. When specific subregions of the NPS were examined, pain-related patterns in the anterior cingulate cortex, bilateral insula, thalamus, secondary sensory cortex and vermis responded to both breathlessness and finger opposition, with all breathlessness results (except the vermis) replicated in an independent study at 3 Tesla. Additionally, the insula, thalamus and secondary somatosensory cortex (S2) were all modulated by the administration of the opioid remifentanil. The activity in these areas may thus provide a general substrate for motivated action within the pain response. In contrast, no conditions positively activated the local NPS pattern in the dorsal posterior insula, an area though to be a critical area for pain perception. Therefore, these results provide new information on the boundary conditions for NPS activation, where a non-zero NPS value is not sufficient to discriminate pain from breathlessness, anticipation of breathlessness, and basic sensorimotor activity. These findings contrast with a number of previous studies that have not found anticipatory activity during anticipated pain (Krishnan et al. 2016; López-Solà et al. 2019). The findings thus suggest that new classifiers, perhaps based on conjunctions of local pattern responses in specific areas, may be required to achieve further specificity. In this regard, the dorsal posterior insula (dpIns) may be a key region, as dpIns (and local NPS pattern in this region) is routinely activated during somatic pain (Geuter et al. 2020), but does not appear to respond to any of the challenges studied here.

### Implications for the understanding of breathlessness

Our findings may also provide insight into the similarities and differences underlying these somatosensory (and often salient) conditions. Current theories regarding the mechanisms and potential treatments for chronic breathlessness often draw heavily on pain models (Parshall et al. 2012; Leupoldt et al. 2009; Lansing et al. 2009), which is understandable considering that they share some phenomenological characteristics. However, with the search for individualised neuro-markers and brain-based treatments for breathlessness becoming an increasing topic of interest (Marlow et al. 2019; Herigstad et al. 2017), it is imperative to attempt to understand what is specific for breathlessness within brain activity and connectivity patterns, rather than over-rely on models created from other conditions.

### Specificity of neural pain signatures

These results help us to understand and explore the current boundaries of an established neural pain signature. While NPS-related activity was significantly activated by non-pain conditions, qualitative pattern differences existed within the regional responses across specific areas. Notably, while sensorimotor areas and the bilateral insula were repeatedly activated by non-painful but somatosensory tasks, the dorsal posterior insula was not positively activated by any of the conditions tested here. The dorsal posterior insula has been frequently implicated as having a critical role in pain perception (Henderson et al. 2010; Brooks et al. 2005; Singer et al. 2004; Ito 1998; Craig 2013; Segerdahl et al. 2015), and may be an essential area in differentiating pain from other salient symptoms. Previous work in both animals (Ito 1998; Craig 2013) and humans (Segerdahl et al. 2015) has determined a subregion of the dorsal posterior insula to be a cortical representation of afferent nociceptive stimuli, and thus it could be considered as an important primary sensory junction for ascending peripheral pain stimuli. Therefore, it is possible that localized patterns of activity in this specific area of the brain may prove more informative for specific determination of painful from non-painful stimuli.

### Neural signatures of motivated actions

While the brain is thought to contain primary cortices dedicated to specific sensory experiences such as vision, audition and touch (Liang et al. 2013; Kwong et al. 1992; Noesselt et al. 2007; Goel et al. 2006), processing of sensory signals does not stop at these junctures. We must de-code these sensory inputs – together with our expectations of the world around us (Seth 2013; Stephan et al. 2016; Van den Bergh et al. 2017; Feldman Barrett & Simmons 2015; Marlow et al. 2019) – to determine what they mean for elements of our health and happiness, and the potential necessity for any further action. Thus, processing these multiple dimensions of perceptual information requires higher cortical involvement and communications beyond primary sensory cortices. While multivariate, brain-wide signatures such as the NPS have been developed to specifically determine the pattern of activity associated with perceptions of somatosensory pain (Wager et al. 2013; Woo, Schmidt, et al. 2017), these complex, salient experiences may not be easily discernable from other threatening perceptions or even simply motivated behaviors in some cases.

Here, we have shown that not only does breathlessness evoke similar patterns of brain activity to that of painful stimuli, but also that anticipating breathlessness and even a simple finger opposition task can both significantly activate the NPS. While the lived experience of these conditions informs us that they are usually easily separable and distinct experiences, they must share common threads within both their nature and activated brain networks. In essence, they all involve the translation of sensory signals to desired motivated behaviors: to avoid the painful stimulus, to overcome the inspiratory resistance (or to prepare for this during an anticipatory period), and to conduct finger opposition movements. When we consider the regional NPS responses to these conditions within the brain, we observe statistical similarities between pain, breathlessness and finger opposition in the thalamus, secondary sensory cortex, bilateral insula and dorsal anterior cingulate cortex. These areas are indeed associated with early sensory processing (thalamus and secondary sensory cortex) (Craig et al. 1994; Ohara & Lenz 2003; Ploner et al. 1999), representations of bodily state (insula) (Singer et al. 2009; Craig 2002; Craig 2009; Craig 2003) and context-specific behaviors towards directed goals (dorsal anterior cingulate) (Holroyd & Yeung 2012), and thus may provide a representative network of sensation-motivated behaviors. However, as anticipation of breathlessness can also induce significant activity in the NPS, it does not appear that the presence of sensory information flow from the periphery is a necessity to activate this blueprint of ‘motivated action’. Rather, the preparatory, future-oriented intent for motivated action may be powerful enough to elicit an NPS-related brain response. Notably, many other salient, motivationally relevant affective conditions have failed to produce NPS activation in previous studies. One possibility for the discrepancy between these studies and the present ones is that many previous comparison conditions involved emotional responses, which appear to engage substantially different brain systems overall from those engaged by pain. Perhaps finger opposition, counterintuitively, produces activity patterns more similar to the NPS because it engages basic motivational, attentional and action processes without the additional different systems engaged during emotion.

There is one important additional caveat. It is unclear from the present results alone whether the degree of activation to breathlessness and its anticipation is comparable to that elicited by somatic pain (Wager et al. 2013), where a quantitative threshold was needed to separate pain from non-painful stimuli, as emotion and non-painful warmth produced relative NPS response differences in the sub-pain-threshold range. Therefore, we cannot know for sure whether the NPS responses observed here are quantitatively strong enough to be classified as “pain” by the original model. Because BOLD signal is not measured in absolute units it remains a challenge to be addressed in the future to compare NPS responses (and other metrics) quantitatively across studies. To further complicate matters, the added signal and statistical power provided across field strengths (such as using 7 Tesla) or between different conditions (such as pain, breathlessness and finger opposition) may overwhelm prescribed magnitude ‘thresholds’ for NPS activity, and thus also need to be considered. While further experimentation including pain, breathing and sensorimotor tasks within one session at the same field strength may shed light on these magnitude differences, the current results do appear to inform us that a simple ‘significant’ activation of these signatures cannot constitute ‘pain’ alone. Moreover, the fact that we observed some NPS activity here in response to non-somatic pain conditions motivates the development and validation of other types of models.

### Conclusions and future directions

So, what do these results mean for the NPS? And for our understanding of breathlessness? Are we chasing the impossible, where a pattern of whole-brain activity can identify pain and pain alone in an individual? And what would the perception of pain become, if the component comprising motivated behavior were removed? We could strive for finer resolutions and better pattern recognition algorithms, with the hope that this specificity exists underneath the noise of functional neuroimaging. Or, with the inherent spatial constraints imposed upon us, and the diversity of brains among us (Gordon et al. 2017), it may be more fruitful to move away from a modular view of the (non-invasively accessible) macro-scale brain, and consider that the existence of a highly specific ‘pain activity network’ may not be achievable given both the importance of cognitive context in shaping pain and the current functional neuroimaging tools (Atlas & Wager 2012; Wiech et al. 2008). That is, somatic conditions such as breathlessness and finger opposition, and even types of anticipatory threat that are sufficiently intense and strongly referred to the body may activate (what has been thought of as) somatic ‘pain’ systems.

Alternatively, we could narrow our initial search to more primary sensory cortices that have repeatedly been associated specifically with pain, such as the dorsal posterior insula (Henderson et al. 2010; Brooks et al. 2005; Singer et al. 2004; Ito 1998; Craig 2013; Segerdahl et al. 2015). These localized patterns could then be combined using rule-based classifiers or combined with brain-wide indicators, and possibly extended and combined with more intricate measures of regional connectivity patterns within dynamic functional networks (Woo et al. 2015). Thus, the present results, alongside animal neuroscience studies showing high specificity of neural populations for particular subtypes of pain and body locations, offer substantial promise for developing pain-specific and breathlessness-specific signature patterns. Understanding both brain activity and connectivity may also provide clues as to the flow of information between primary sensory cortices and higher cognitive and limbic structures, and may thus offer the required specificity to help us identify pain in those who cannot express it for themselves.

## Acknowledgements

Olivia Harrison (née Faull) is a Marie Skłodowska-Curie Postdoctoral Fellow, supported by the European Union’s Horizon 2020 research and innovation programme under the Grant Agreement No 793580. This work was supported in part by NIH grants R01DA035484, 2R01MH076136 and R01DA027794 (TDW). Kyle Pattinson was supported by the JABBS Foundation, the Dunhill Medical Trust, and the NIHR Biomedical Research Centre based at Oxford University Hospitals NHS Trust and The University of Oxford. This article is based upon work funded by a Medical Research Council Clinician Scientist Fellowship (grant number G0802826) awarded to K.T.S.P.

## Supplementary Material

**Supplementary Table 1:**
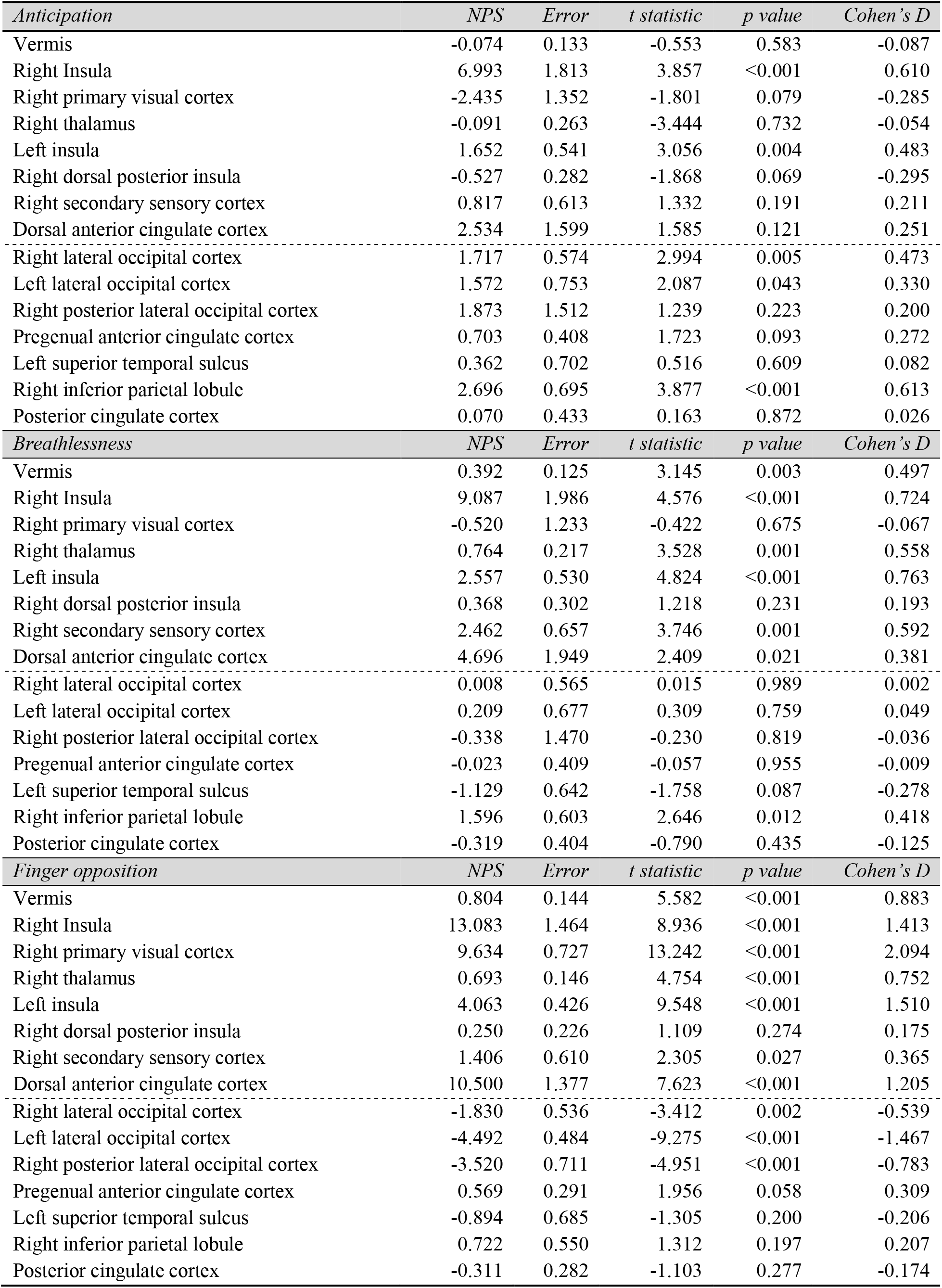
NPS subregion analyses for 7 Tesla data contrasts of interest. Positive regions above the dotted line, negative regions below.

| *Anticipation* | *NPS* | *Error* | *t statistic* | *p value* | *Cohen's D* |
| --- | --- | --- | --- | --- | --- |
| Vermis | -0.074 | 0.133 | -0.553 | 0.583 | -0.087 |
| Right Insula | 6.993 | 1.813 | 3.857 | <0.001 | 0.610 |
| Right primary visual cortex | -2.435 | 1.352 | -1.801 | 0.079 | -0.285 |
| Right thalamus | -0.091 | 0.263 | -3.444 | 0.732 | -0.054 |
| Left insula | 1.652 | 0.541 | 3.056 | 0.004 | 0.483 |
| Right dorsal posterior insula | -0.527 | 0.282 | -1.868 | 0.069 | -0.295 |
| Right secondary sensory cortex | 0.817 | 0.613 | 1.332 | 0.191 | 0.211 |
| Dorsal anterior cingulate cortex | 2.534 | 1.599 | 1.585 | 0.121 | 0.251 |
| Right lateral occipital cortex | 1.717 | 0.574 | 2.994 | 0.005 | 0.473 |
| Left lateral occipital cortex | 1.572 | 0.753 | 2.087 | 0.043 | 0.330 |
| Right posterior lateral occipital cortex | 1.873 | 1.512 | 1.239 | 0.223 | 0.200 |
| Pregenual anterior cingulate cortex | 0.703 | 0.408 | 1.723 | 0.093 | 0.272 |
| Left superior temporal sulcus | 0.362 | 0.702 | 0.516 | 0.609 | 0.082 |
| Right inferior parietal lobule | 2.696 | 0.695 | 3.877 | <0.001 | 0.613 |
| Posterior cingulate cortex | 0.070 | 0.433 | 0.163 | 0.872 | 0.026 |
| *Breathlessness* | *NPS* | *Error* | *t statistic* | *p value* | *Cohen's D* |
| Vermis | 0.392 | 0.125 | 3.145 | 0.003 | 0.497 |
| Right Insula | 9.087 | 1.986 | 4.576 | <0.001 | 0.724 |
| Right primary visual cortex | -0.520 | 1.233 | -0.422 | 0.675 | -0.067 |
| Right thalamus | 0.764 | 0.217 | 3.528 | 0.001 | 0.558 |
| Left insula | 2.557 | 0.530 | 4.824 | <0.001 | 0.763 |
| Right dorsal posterior insula | 0.368 | 0.302 | 1.218 | 0.231 | 0.193 |
| Right secondary sensory cortex | 2.462 | 0.657 | 3.746 | 0.001 | 0.592 |
| Dorsal anterior cingulate cortex | 4.696 | 1.949 | 2.409 | 0.021 | 0.381 |
| Right lateral occipital cortex | 0.008 | 0.565 | 0.015 | 0.989 | 0.002 |
| Left lateral occipital cortex | 0.209 | 0.677 | 0.309 | 0.759 | 0.049 |
| Right posterior lateral occipital cortex | -0.338 | 1.470 | -0.230 | 0.819 | -0.036 |
| Pregenual anterior cingulate cortex | -0.023 | 0.409 | -0.057 | 0.955 | -0.009 |
| Left superior temporal sulcus | -1.129 | 0.642 | -1.758 | 0.087 | -0.278 |
| Right inferior parietal lobule | 1.596 | 0.603 | 2.646 | 0.012 | 0.418 |
| Posterior cingulate cortex | -0.319 | 0.404 | -0.790 | 0.435 | -0.125 |
| *Finger opposition* | *NPS* | *Error* | *t statistic* | *p value* | *Cohen's D* |
| Vermis | 0.804 | 0.144 | 5.582 | <0.001 | 0.883 |
| Right Insula | 13.083 | 1.464 | 8.936 | <0.001 | 1.413 |
| Right primary visual cortex | 9.634 | 0.727 | 13.242 | <0.001 | 2.094 |
| Right thalamus | 0.693 | 0.146 | 4.754 | <0.001 | 0.752 |
| Left insula | 4.063 | 0.426 | 9.548 | <0.001 | 1.510 |
| Right dorsal posterior insula | 0.250 | 0.226 | 1.109 | 0.274 | 0.175 |
| Right secondary sensory cortex | 1.406 | 0.610 | 2.305 | 0.027 | 0.365 |
| Dorsal anterior cingulate cortex | 10.500 | 1.377 | 7.623 | <0.001 | 1.205 |
| Right lateral occipital cortex | -1.830 | 0.536 | -3.412 | 0.002 | -0.539 |
| Left lateral occipital cortex | -4.492 | 0.484 | -9.275 | <0.001 | -1.467 |
| Right posterior lateral occipital cortex | -3.520 | 0.711 | -4.951 | <0.001 | -0.783 |
| Pregenual anterior cingulate cortex | 0.569 | 0.291 | 1.956 | 0.058 | 0.309 |
| Left superior temporal sulcus | -0.894 | 0.685 | -1.305 | 0.200 | -0.206 |
| Right inferior parietal lobule | 0.722 | 0.550 | 1.312 | 0.197 | 0.207 |
| Posterior cingulate cortex | -0.311 | 0.282 | -1.103 | 0.277 | -0.174 |

**Supplementary Table 2:**
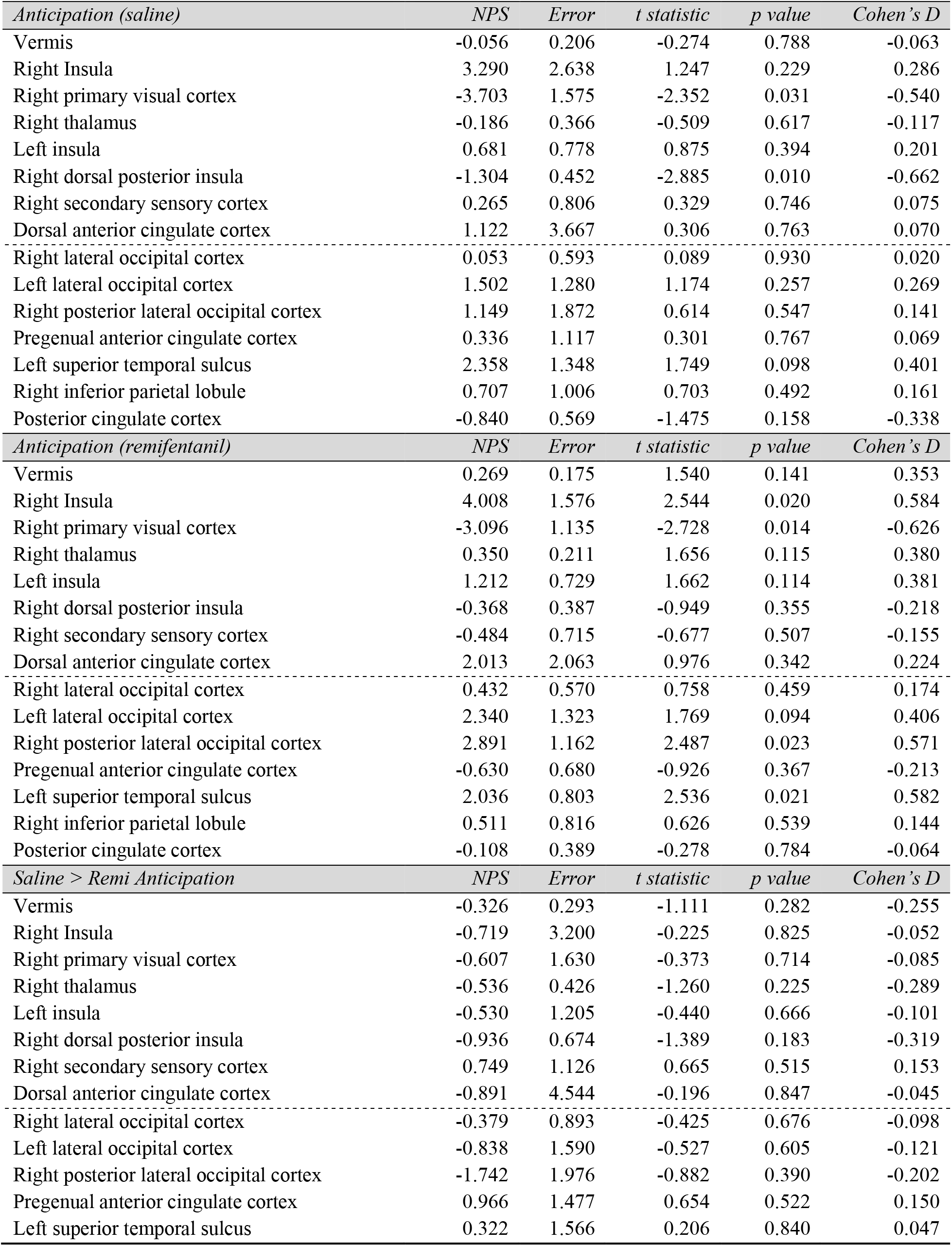
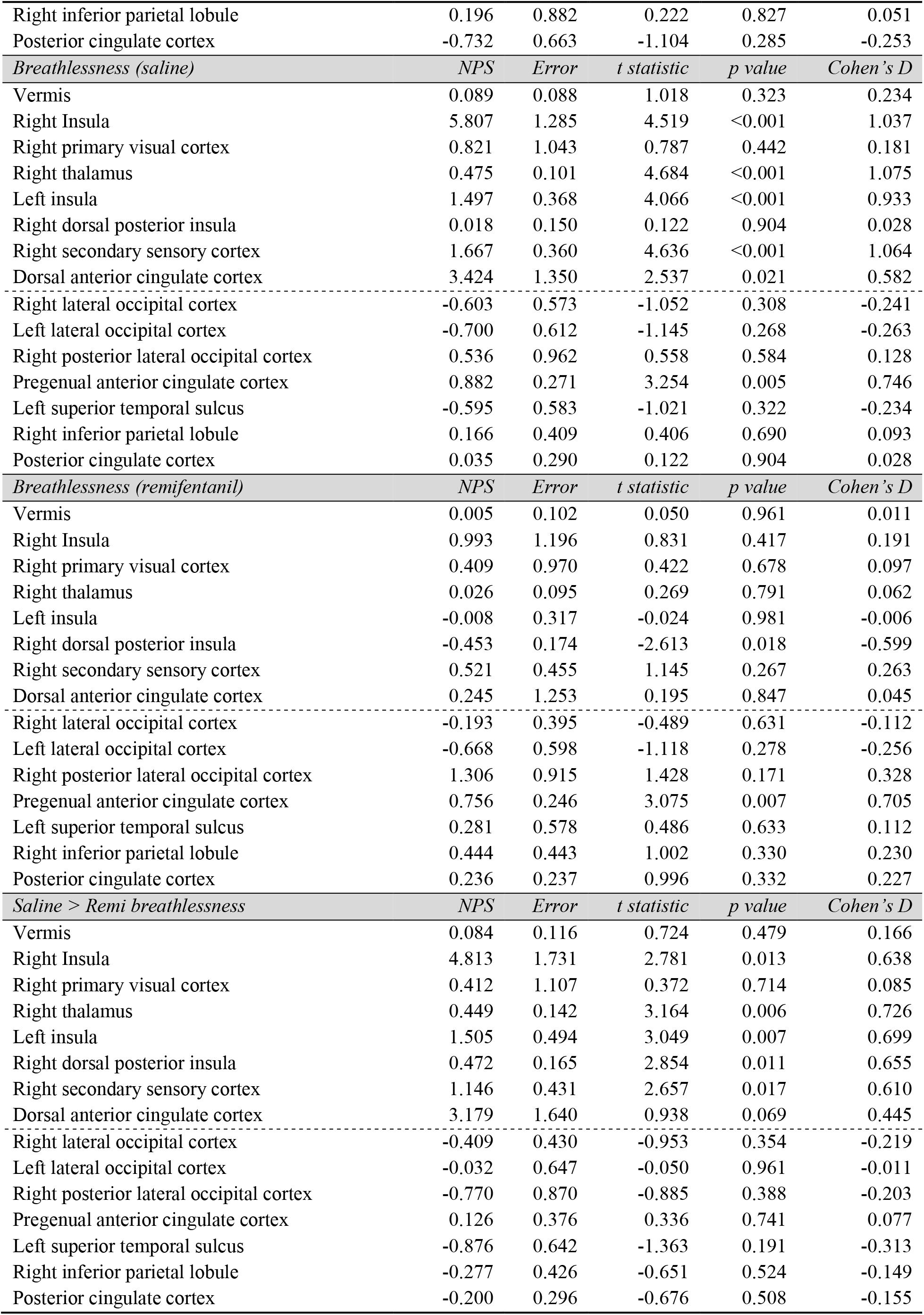
NPS subregion analyses for 3 Tesla data contrasts of interest. Positive regions above the dotted line, negative regions below.

| *Anticipation (saline)* | *NPS* | *Error* | *t statistic* | *p value* | *Cohen's D* |
| --- | --- | --- | --- | --- | --- |
| Vermis | -0.056 | 0.206 | -0.274 | 0.788 | -0.063 |
| Right Insula | 3.290 | 2.638 | 1.247 | 0.229 | 0.286 |
| Right primary visual cortex | -3.703 | 1.575 | -2.352 | 0.031 | -0.540 |
| Right thalamus | -0.186 | 0.366 | -0.509 | 0.617 | -0.117 |
| Left insula | 0.681 | 0.778 | 0.875 | 0.394 | 0.201 |
| Right dorsal posterior insula | -1.304 | 0.452 | -2.885 | 0.010 | -0.662 |
| Right secondary sensory cortex | 0.265 | 0.806 | 0.329 | 0.746 | 0.075 |
| Dorsal anterior cingulate cortex | 1.122 | 3.667 | 0.306 | 0.763 | 0.070 |
| Right lateral occipital cortex | 0.053 | 0.593 | 0.089 | 0.930 | 0.020 |
| Left lateral occipital cortex | 1.502 | 1.280 | 1.174 | 0.257 | 0.269 |
| Right posterior lateral occipital cortex | 1.149 | 1.872 | 0.614 | 0.547 | 0.141 |
| Pregenual anterior cingulate cortex | 0.336 | 1.117 | 0.301 | 0.767 | 0.069 |
| Left superior temporal sulcus | 2.358 | 1.348 | 1.749 | 0.098 | 0.401 |
| Right inferior parietal lobule | 0.707 | 1.006 | 0.703 | 0.492 | 0.161 |
| Posterior cingulate cortex | -0.840 | 0.569 | -1.475 | 0.158 | -0.338 |
| *Anticipation (remifentanyl)* | *NPS* | *Error* | *t statistic* | *p value* | *Cohen's D* |
| Vermis | 0.269 | 0.175 | 1.540 | 0.141 | 0.353 |
| Right Insula | 4.008 | 1.576 | 2.544 | 0.020 | 0.584 |
| Right primary visual cortex | -3.096 | 1.135 | -2.728 | 0.014 | -0.626 |
| Right thalamus | 0.350 | 0.211 | 1.656 | 0.115 | 0.380 |
| Left insula | 1.212 | 0.729 | 1.662 | 0.114 | 0.381 |
| Right dorsal posterior insula | -0.368 | 0.387 | -0.949 | 0.355 | -0.218 |
| Right secondary sensory cortex | -0.484 | 0.715 | -0.677 | 0.507 | -0.155 |
| Dorsal anterior cingulate cortex | 2.013 | 2.063 | 0.976 | 0.342 | 0.224 |
| Right lateral occipital cortex | 0.432 | 0.570 | 0.758 | 0.459 | 0.174 |
| Left lateral occipital cortex | 2.340 | 1.323 | 1.769 | 0.094 | 0.406 |
| Right posterior lateral occipital cortex | 2.891 | 1.162 | 2.487 | 0.023 | 0.571 |
| Pregenual anterior cingulate cortex | -0.630 | 0.680 | -0.926 | 0.367 | -0.213 |
| Left superior temporal sulcus | 2.036 | 0.803 | 2.536 | 0.021 | 0.582 |
| Right inferior parietal lobule | 0.511 | 0.816 | 0.626 | 0.539 | 0.144 |
| Posterior cingulate cortex | -0.108 | 0.389 | -0.278 | 0.784 | -0.064 |
| *Saline > Remi Anticipation* | *NPS* | *Error* | *t statistic* | *p value* | *Cohen's D* |
| Vermis | -0.326 | 0.293 | -1.111 | 0.282 | -0.255 |
| Right Insula | -0.719 | 3.200 | -0.225 | 0.825 | -0.052 |
| Right primary visual cortex | -0.607 | 1.630 | -0.373 | 0.714 | -0.085 |
| Right thalamus | -0.536 | 0.426 | -1.260 | 0.225 | -0.289 |
| Left insula | -0.530 | 1.205 | -0.440 | 0.666 | -0.101 |
| Right dorsal posterior insula | -0.936 | 0.674 | -1.389 | 0.183 | -0.319 |
| Right secondary sensory cortex | 0.749 | 1.126 | 0.665 | 0.515 | 0.153 |
| Dorsal anterior cingulate cortex | -0.891 | 4.544 | -0.196 | 0.847 | -0.045 |
| Right lateral occipital cortex | -0.379 | 0.893 | -0.425 | 0.676 | -0.098 |
| Left lateral occipital cortex | -0.838 | 1.590 | -0.527 | 0.605 | -0.121 |
| Right posterior lateral occipital cortex | -1.742 | 1.976 | -0.882 | 0.390 | -0.202 |
| Pregenual anterior cingulate cortex | 0.966 | 1.477 | 0.654 | 0.522 | 0.150 |
| Left superior temporal sulcus | 0.322 | 1.566 | 0.206 | 0.840 | 0.047 |
| Right inferior parietal lobule | 0.196 | 0.882 | 0.222 | 0.827 | 0.051 |
| Posterior cingulate cortex | -0.732 | 0.663 | -1.104 | 0.285 | -0.253 |
| *Breathlessness (saline)* | *NPS* | *Error* | *t statistic* | *p value* | *Cohen's D* |
| Vermis | 0.089 | 0.088 | 1.018 | 0.323 | 0.234 |
| Right Insula | 5.807 | 1.285 | 4.519 | <0.001 | 1.037 |
| Right primary visual cortex | 0.821 | 1.043 | 0.787 | 0.442 | 0.181 |
| Right thalamus | 0.475 | 0.101 | 4.684 | <0.001 | 1.075 |
| Left insula | 1.497 | 0.368 | 4.066 | <0.001 | 0.933 |
| Right dorsal posterior insula | 0.018 | 0.150 | 0.122 | 0.904 | 0.028 |
| Right secondary sensory cortex | 1.667 | 0.360 | 4.636 | <0.001 | 1.064 |
| Dorsal anterior cingulate cortex | 3.424 | 1.350 | 2.537 | 0.021 | 0.582 |
| Right lateral occipital cortex | -0.603 | 0.573 | -1.052 | 0.308 | -0.241 |
| Left lateral occipital cortex | -0.700 | 0.612 | -1.145 | 0.268 | -0.263 |
| Right posterior lateral occipital cortex | 0.536 | 0.962 | 0.558 | 0.584 | 0.128 |
| Pregenual anterior cingulate cortex | 0.882 | 0.271 | 3.254 | 0.005 | 0.746 |
| Left superior temporal sulcus | -0.595 | 0.583 | -1.021 | 0.322 | -0.234 |
| Right inferior parietal lobule | 0.166 | 0.409 | 0.406 | 0.690 | 0.093 |
| Posterior cingulate cortex | 0.035 | 0.290 | 0.122 | 0.904 | 0.028 |
| *Breathlessness (remifentanyl)* | *NPS* | *Error* | *t statistic* | *p value* | *Cohen's D* |
| Vermis | 0.005 | 0.102 | 0.050 | 0.961 | 0.011 |
| Right Insula | 0.993 | 1.196 | 0.831 | 0.417 | 0.191 |
| Right primary visual cortex | 0.409 | 0.970 | 0.422 | 0.678 | 0.097 |
| Right thalamus | 0.026 | 0.095 | 0.269 | 0.791 | 0.062 |
| Left insula | -0.008 | 0.317 | -0.024 | 0.981 | -0.006 |
| Right dorsal posterior insula | -0.453 | 0.174 | -2.613 | 0.018 | -0.599 |
| Right secondary sensory cortex | 0.521 | 0.455 | 1.145 | 0.267 | 0.263 |
| Dorsal anterior cingulate cortex | 0.245 | 1.253 | 0.195 | 0.847 | 0.045 |
| Right lateral occipital cortex | -0.193 | 0.395 | -0.489 | 0.631 | -0.112 |
| Left lateral occipital cortex | -0.668 | 0.598 | -1.118 | 0.278 | -0.256 |
| Right posterior lateral occipital cortex | 1.306 | 0.915 | 1.428 | 0.171 | 0.328 |
| Pregenual anterior cingulate cortex | 0.756 | 0.246 | 3.075 | 0.007 | 0.705 |
| Left superior temporal sulcus | 0.281 | 0.578 | 0.486 | 0.633 | 0.112 |
| Right inferior parietal lobule | 0.444 | 0.443 | 1.002 | 0.330 | 0.230 |
| Posterior cingulate cortex | 0.236 | 0.237 | 0.996 | 0.332 | 0.227 |
| *Saline > Remi breathlessness* | *NPS* | *Error* | *t statistic* | *p value* | *Cohen's D* |
| Vermis | 0.084 | 0.116 | 0.724 | 0.479 | 0.166 |
| Right Insula | 4.813 | 1.731 | 2.781 | 0.013 | 0.638 |
| Right primary visual cortex | 0.412 | 1.107 | 0.372 | 0.714 | 0.085 |
| Right thalamus | 0.449 | 0.142 | 3.164 | 0.006 | 0.726 |
| Left insula | 1.505 | 0.494 | 3.049 | 0.007 | 0.699 |
| Right dorsal posterior insula | 0.472 | 0.165 | 2.854 | 0.011 | 0.655 |
| Right secondary sensory cortex | 1.146 | 0.431 | 2.657 | 0.017 | 0.610 |
| Dorsal anterior cingulate cortex | 3.179 | 1.640 | 0.938 | 0.069 | 0.445 |
| Right lateral occipital cortex | -0.409 | 0.430 | -0.953 | 0.354 | -0.219 |
| Left lateral occipital cortex | -0.032 | 0.647 | -0.050 | 0.961 | -0.011 |
| Right posterior lateral occipital cortex | -0.770 | 0.870 | -0.885 | 0.388 | -0.203 |
| Pregenual anterior cingulate cortex | 0.126 | 0.376 | 0.336 | 0.741 | 0.077 |
| Left superior temporal sulcus | -0.876 | 0.642 | -1.363 | 0.191 | -0.313 |
| Right inferior parietal lobule | -0.277 | 0.426 | -0.651 | 0.524 | -0.149 |
| Posterior cingulate cortex | -0.200 | 0.296 | -0.676 | 0.508 | -0.155 |

## Supplementary Figures

**Supplementary Figure 1.**
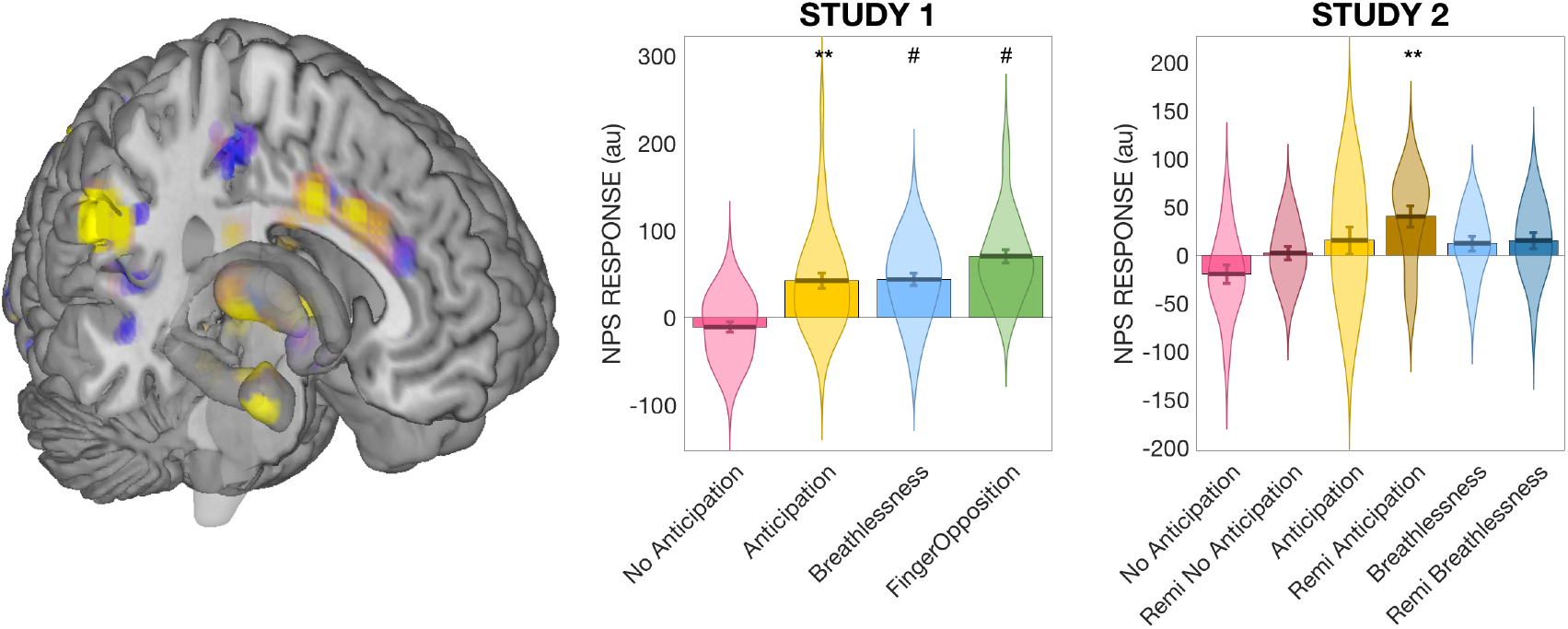
Overall NPS activity in all conditions for the two datasets. Left: Three-dimensional representation of some of the core regions of the NPS. ** Significantly different from zero at p < 0.01; # Significantly different from zero at q < 0.05 (FDR corrected).

**Supplementary Figure 2.**
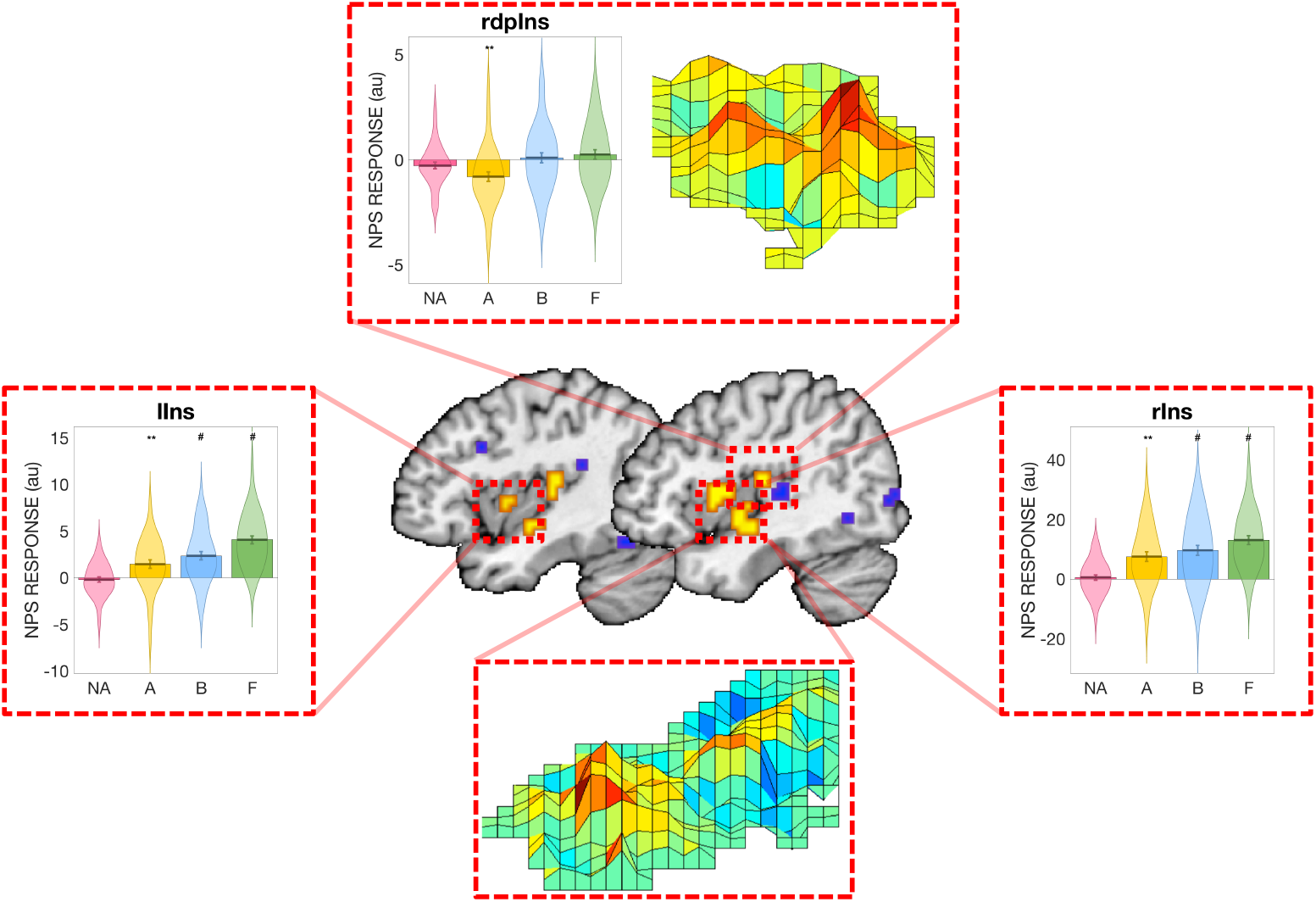
Regional NPS activity in the insula for the no-anticipation, anticipation, breathlessness and finger opposition conditions from Study 1. Robust statistical activity is observed in the bilateral insula (labelled lIns and rIns) for all except the no-anticipation condition, while no significant positive activity is observed in the right dorsal posterior insula (rdpIns). Abbreviations: NA, No anticipation; A, Anticipation; B, Breathlessness; F, Finger opposition. ** Significantly different from zero at p < 0.01; # Significantly different from zero at q < 0.05 (FDR corrected).

**Supplementary Figure 3.**
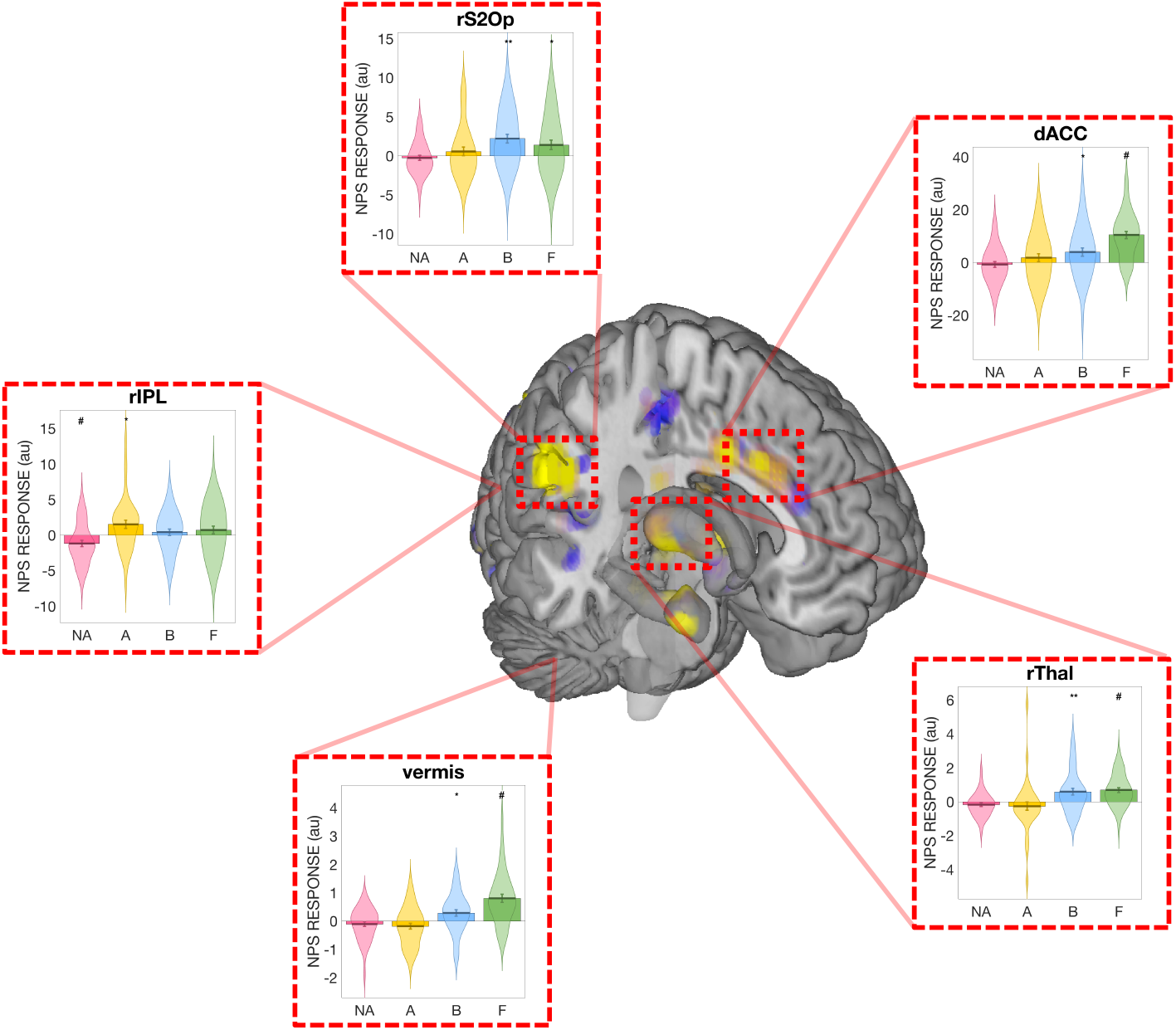
Regional NPS activity subregions of the NPS for the no-anticipation, anticipation, breathlessness and finger opposition conditions from Study 1. Abbreviations: dACC, dorsal anterior cingulate cortex; rThal, right thalamus; rS2Op, right secondary somatosensory cortex / operculum; rIPL, right inferior parietal lobule; NA, No anticipation; A, Anticipation; B, Breathlessness; F, Finger opposition. * Significantly different from zero at p < 0.05; ** Significantly different from zero at p < 0.01; # Significantly different from zero at q < 0.05 (FDR corrected).

**Supplementary Figure 4.**
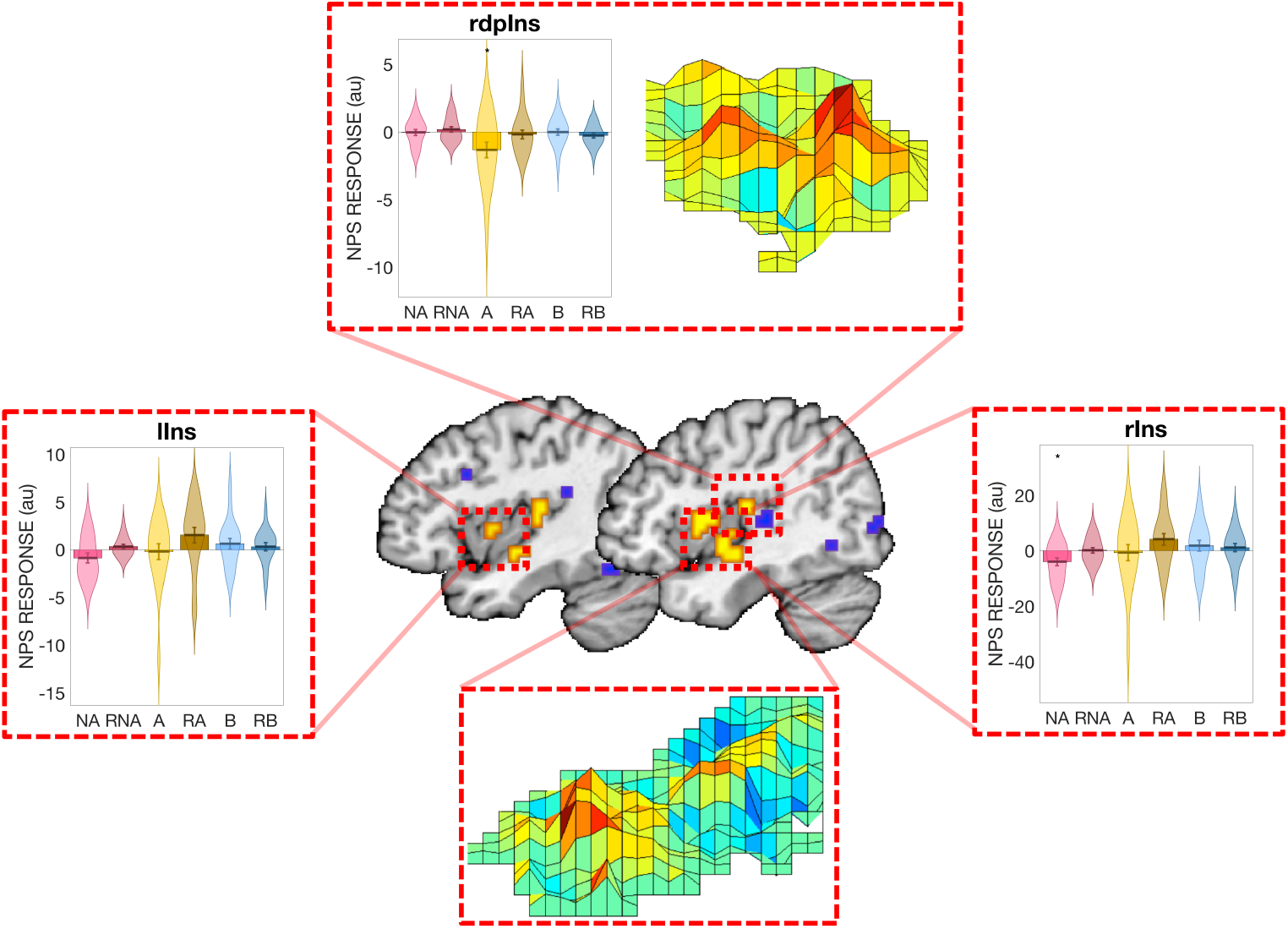
Regional NPS activity in the insula for the no-anticipation, anticipation and breathlessness conditions during both saline and remifentanil administration from Study 2. Abbreviations: rIns, right insula; lIns, left insula; rdpIns, right dorsal posterior insula; NA, No anticipation; A, Anticipation (saline); RA, Remifentanil anticipation; B, Breathlessness (saline); RB, Remifentanil breathlessness. ** Significantly different from zero at p < 0.01; # Significantly different from zero at q < 0.05 (FDR corrected).

**Supplementary Figure 5.**
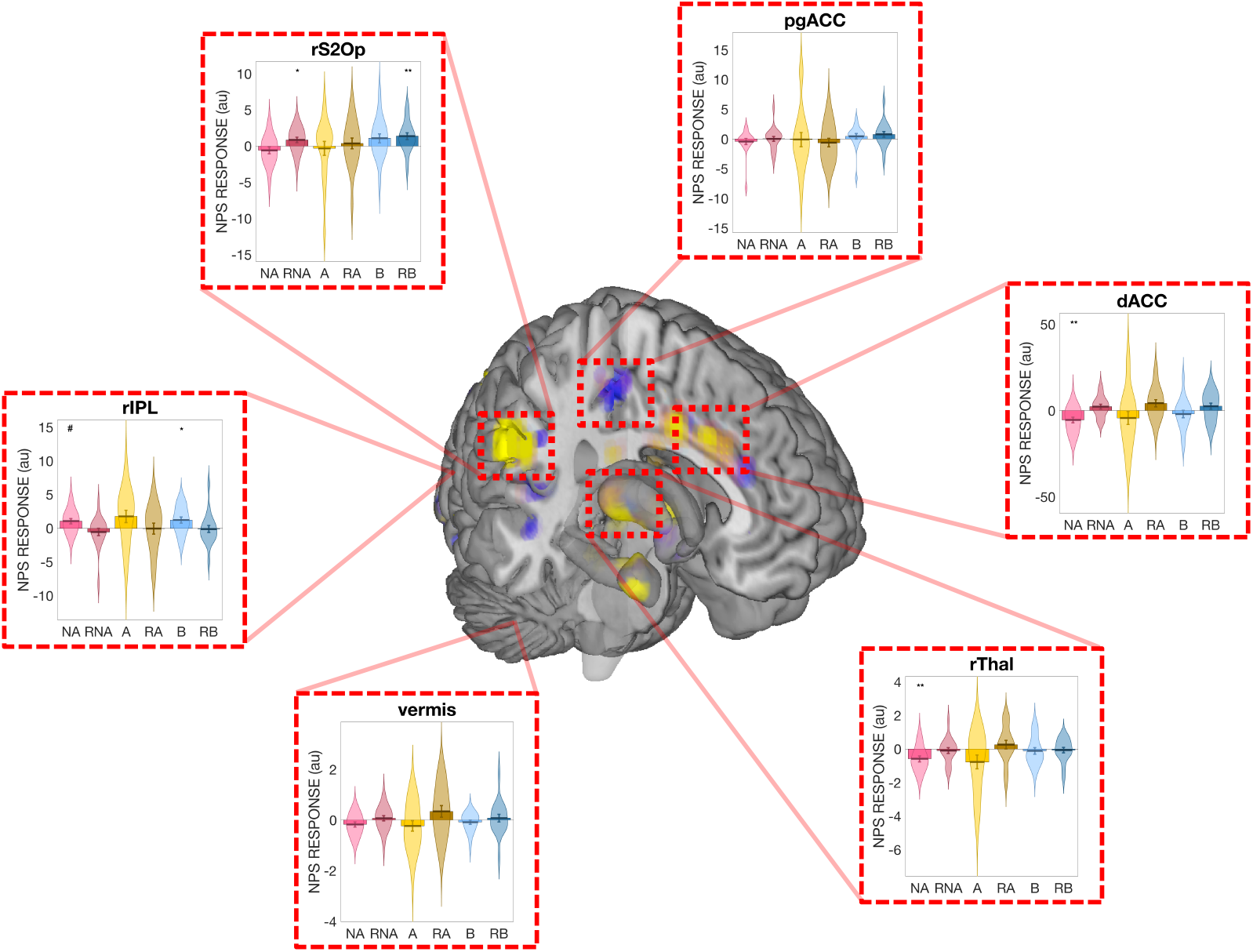
Regional NPS activity subregions of the NPS for the no-anticipation, anticipation and breathlessness conditions during both saline and remifentanil administration from Study 2. Abbreviations: dACC, dorsal anterior cingulate cortex; pgACC, pregenual anterior cingulate cortex; rThal right thalamus; rS2Op, right secondary somatosensory cortex / operculum; rIPL, right inferior parietal lobule; A, Anticipation contrast (saline); RA, Remifentanil anticipation contrast; B, Breathlessness contrast (saline); RB, Remifentanil breathlessness contrast. * Significantly different from zero at p < 0.05; ** Significantly different from zero at p < 0.01; # Significantly different from zero at q < 0.05 (FDR corrected).

